# SnoVault and encodeD: A novel object-based storage system and applications to ENCODE metadata

**DOI:** 10.1101/044578

**Authors:** Benjamin C. Hitz, Laurence D. Rowe, Nikhil R. Podduturi, David I. Glick, Ulugbek K. Baymuradov, Venkat S. Malladi, Esther T. Chan, Jean M. Davidson, Idan Gabdank, Aditi K. Narayanan, Kathrina C. Onate, Jason Hilton, Marcus C. Ho, Brian T. Lee, Stuart R. Miyasato, Timothy R. Dreszer, Cricket A. Sloan, J. Seth Strattan, Forrest Y. Tanaka, Eurie L. Hong, J. Michael Cherry

## Abstract

The Encyclopedia of DNA elements (ENCODE) project is an ongoing collaborative effort[1–6] to create a comprehensive catalog of functional elements initiated shortly after the completion of the Human Genome Project[7][1]. The current database exceeds 6500 experiments across more than 450 cell lines and tissues using a wide array of experimental techniques to study the chromatin structure, regulatory and transcriptional landscape of the *H. sapiens* and *M. musculus* genomes. All ENCODE experimental data, metadata, and associated computational analyses are submitted to the ENCODE Data Coordination Center (DCC) for validation, tracking, storage, unified processing, and distribution to community resources and the scientific community. As the volume of data increases, the identification and organization of experimental details becomes increasingly intricate and demands careful curation. The ENCODE DCC[8–10] has created a general purpose software system, known as SnoVault, that supports metadata and file submission, a database used for metadata storage, web pages for displaying the metadata and a robust API for querying the metadata. The software is fully open-source, code and installation instructions can be found at: http://github.com/ENCODE-DCC/snovault/ (for the generic database) and http://github.com/ENCODE-DCC/encoded/ to store genomic data in the manner of ENCODE. The core database engine, SnoVault (which is completely independent of ENCODE, genomic data, or bioinformatic data) has been released as a separate Python package.

**Database URL:** https://www.encodeproject.org/

## Introduction

The Encyclopedia of DNA Elements (ENCODE) project (https://www.encodeproject.org/) is an international consortium with a goal of annotating regions of the genome [1–6]. The Data Coordination Center (DCC) is charged with validating, tracking, storing, processing, visualizing and distributing these data files and their metadata to the scientific community. During 6 years of the pilot and initial scale-up phase, the project surveyed the landscape of the *H. sapiens* and *M. musculus* genomes using over 20 high-throughput genomic assays in more than 350 different cell and tissue types, resulting in over 3000 datasets. ENCODE employs a wide range of genomics techniques[11, 12]; [13]. To determine the transcribed elements including mRNAs, the GENCODE (http://www.gencodegenes.org) project employs a team of biocurators who synthesize data from a wide range of assays to build accurate gene models that address the full complexity of vertebrate transcription including alternative promoters, alternative splicing, and RNA editing[14]. These assays include sequencing selected RNAs for promoter characterization, next-generation RNA sequencing and assembly methods for detecting alternative splicing and for quantizing the level of RNA in different cellular contexts[15]. To determine transcriptional regulatory regions ENCODE uses DNase I [15, 16]and nucleosome positioning assays to locate regions of the chromosome accessible to regulatory elements, DNA methylation assays and ChIP-seq of modified histones to define the overall chromatin architecture[17][18], and ChIP-seq of transcription factors to determine the players involved in the regulatory interactions[19]. To determine which regulatory elements interact with each other, ENCODE uses ChIA-PE T[19, 20]and Hi-C[21]. To identify regulatory elements that operate on the RNA rather than the DNA level, ENCODE employs RIP-seq[22] as well as a variety of computational assays[23, 24];[25][23, 24][26]. Over the current phase of ENCODE that began in 2012, the diversity and volume of data has increased as new genomic assays are added to the project In addition to data from experiments performed in diversity of biological samples, data from additional species *(D. melanogaster* and *C. elegans)* [27] and other projects have been incorporated. Experimental data are validated and analyzed using new methods, and higher level analyses have been included to form a true Encyclopedia of genomic elements. The DCC currently houses experimental data and metadata from ENCODE3 (the current ENCODE), modENCODE[28], ENCODE2[29](the phase of ENCODE ending in 2012), mouse-ENCODE[29, 30], Roadmap for Epigenomics Mapping (REMC)[29–31], modERN (fly and worm experiments associated with ENCODE3) and Genomics for Gene Regulation (GGR) (https://www.genome.gov/27561317). These contributions are summarized on the ENCODE Portal at:https://www.encodeproiect.org/about/data-access/.

The primary goals of the DCC are to track and compile the experimental metadata for each experiment produced by the consortium. This metadata is an absolutely critical resource both for internal consortium metrics and bookkeeping and to provide the best possible user experience for scientists and educators wishing to make sense of such a large corpus of datafiles, experiments, and outputs[8–10].

One of the challenges in this field is keeping a flexible data model without sacrificing data continuity and integrity. To this end, the SnoVault system includes an infrastructure to build (quality or validity) data audits and reports to maintain data integrity. To resolve these conflicting design criteria, we have developed a hybrid relational/object database using JSON, JSON-LD (http://json-ld.org/), JSON-SCHEMA (http://json-schema.org/), and PostgreSQL (http://www.postgresql.org). For efficient searching and display, the data are denormalized and indexed in ElasticSearch (http://www.elasticsearch.org/). These software components are wrapped in a Python Pyramid web framework application which executes the business logic of the application and provides the RESTful web API. The website itself, also called the “front-end” is constructed using the ReactJS framework (https://facebook.github.io/react/).

## A Hybrid Relational-Object Data Store

Traditionally, biological databases have been implemented using a relational database model. Relational database software, both open-source (e.g., MySQL or PostgreSQL) and commercial (e.g. Oracle), is ubiquitous on all current hardware. It is a robust system for maintaining transactional integrity and keeping concepts normalized (storing each data only once, and connecting them via foreign-key relations). In the SnoVault system, each major category of metadata is a JSON object or “document” typically corresponding to an experimental component of a genomic assay, such as an RNA-seq or ChIP-seq assay. For example, there are JSON objects that represent a specific human donor, which can be linked to 1 or more biological samples (biosamples) from that donor as well as objects that represent key reagents in an assay such as an antibody lot that is used in a ChIP-seq assay or an shRNA used to knockdown a gene target. In addition, objects represent the files and computational analyses required to generate these files. In addition to objects that represent experimental components and reagents of a genomic assay, there are objects that allow the grouping of these objects to represent a single biological replicate and a grouping of biological replicates to represent a replicated assay.

These JSON objects are stored in our transactional implementation of an document store in PostgreSQL. This low-level database is analogous to a commercial or OSS object/document store like MongoDB (http://www.mongodb.com) or couchDB (http://couchdb.apache.org) with additional transactional and rollback capabilities. The other critical features we have added to “classical” object storage methods are JSON-SCHEMA and JSON-LD. JSON-SCHEMA is simply a way to template JSON objects to control the allowed fields in each object and their allowed values in each field. There are separate schema files for each object type, which are analogous to tables in a traditional relational database. JSON-LD is a data standards format that provides a unified and straightforward way of linking JSON objects via pointers or “links” (analogous to foreign keys in a traditional relational database). These links are used in the metadata storage in a manner analogous to foreign key relationships in a traditional relational model. These links allow our software to “upgrade” objects to a new version of a schema when the schema changes. Our upgrading system allows us to return objects that are compatible with the most current schema without necessarily reindexing them in ElasticSearch; they are permanently upgraded when the objects are PUT or PATCHed. Herein lies the critical advance we have made - JSON-LD and JSON-SCHEMA allow us to strike a balance between the flexibility of an unstructured object database and the data integrity and normalization features of a relational database.

To date, the encodeD system (built on SnoVault) contains 75 object types that are able to handle over 40 different classes and flavors of genomic assays. The addition of new assays during the course of the ENCODE project has resulted in minimal changes that often include the addition of a new experimental properties in an existing object or a new object to allow new relationships to be created among existing objects. The flexibility of the SnoVault system allows the addition of these new properties or objects with minimal intervention from software engineers. Data wranglers or other data scientists are perfectly capable of extending or modifying the JSON-SCHEMA model, with only a small amount of training.

We us the open-source python web framework Pyramid (http://www.pylonsproject.org/) as the base python layer of our back end. Pyramid’s excellent buildout system (similar in principle to “make”) allows us to manage our dependencies for python, node.js, and ruby and makes installation on an arbitrary UNIX machine very straight forward.

One of the features of ENCODE DCC website and RESTful API is the ability to set permissions (view, edit, or delete) on a single object independent of the objects that might refer to or embed it. Data that are released are publicly viewable without needing a login at all, while metadata that is still incomplete or in progress is available to view by consortium members. Viewing permissions are defined by a combination of the release status of an object. Editing permissions are defined by a combination of the release status of an object and a membership in a user group. An “admin” user group has permission to edit all objects, regardless of release status. Individual users may belong to specific groups, as defined by labs, to allow editing of data that are generated by that lab. An individual user may be authorized to edit only data for 1 lab or as many labs as appropriate. These user groups are only allowed to edit data that is not released. In this way, we distinguish between “admin” level users (such as an ENCODE data wrangler) who have full rights to all data; a “submitter” level user (someone from a production lab) who can modify the objects of a lab that they submit for; a “consortium” user, who can view all submitted data, and a “base” user, who can only view data that is released. Pyramid has excellent support for detailed “Action Control Lists” (ACLs) which were utilized in our application. The ability to edit and view can be customized by defining the set of permissions that are granted to the specific group. Specifically, submitters from the ENCODE and GGR projects are distinguished without our system; members of those groups may neither view or edit unreleased data from another group.

## Indexing and Performance Optimization

One of the challenges in web database development is to create a system that optimized both for WRITE access and storage and for READ access. Storing data requires transactional integrity, flexibility of schema and some level of denormalization. Our JSON-SCHEMA document model using links to other objects for references fulfills this goal, but retrieving complex linked objects using multiple GETs for each linked object (and so on down through the object tree) is very inefficient. We take the classical approach of indexing denormalized data that we need to retrieve rapidly, but add some modern twists. Each object or document in our system has a uuid that acts similar to a primary key in a relational database. Using JSON-LD, we reference other objects by their object type and uuid, in the parlance of JSON-LD these are called links. Lists of objects or more complex data structures can be used as well. Objects can be returned from the RESTful API either with or without embedded linked objects using a URL query parameter (frame=object, frame=embedded, frame=raw or frame=page). An embedded JSON-LD object is one in which it’s links have been substituted with “copies” (or denormalization) of the linked objects (often just a subsection of the linked objects; the details are specified in the python code for the views of each object type).

To make the database performant, and to enable rapid and faceted searching of the metadata database implemented in encodeD, we index all frames of all objects in ElasticSearch (https://www.elastic.co/products/elasticsearch). ElasticSearch is a robust search and live indexing wrapper based around Apache Lucene (https://lucene.apache.org/core/). As new objects are added to the database via POST or modified via PATCH, they are indexed in real time by the indexer process.

The real-time indexing occurs according to the following rubric:

- When rendering a response, we record the set of embedded_uuids and linked_uuids used.
- The embedded_uuids are those objects embedded into the response or whose properties have been consulted in rendering of the response.
- Any change to one of these objects should cause an invalidation.
- Linked_uuids are the objects linked to in the response. Only changes to their URL (i.e, the canonical name of the link) need trigger an invalidation.
- When modifying objects, event subscribers keep track of which objects are updated and their resource paths before and after the modification. This is used to record the set of `updated_uuids` and `renamed_uuids` in the transaction log.
- The indexer process listens for notifications of new transactions.
- With the union of updated_uuids and union of renamed_uuids across each transaction in the log since its last indexing run, the indexer performs a search for all objects where embedded_uuids intersect with the updated_uuids or linked_uuids intersect with the renamed_uuids.
- The result is the set of invalidated objects which must be reindexed in addition to those that were modified (recorded in updated_uuids.)

## RESTful API

Programmatic interaction with the ENCODE DCC metadata database is typically done through scripts written and executed on a local computer. These scripts interact with the database through an industry-standard, Hypertext-Transfer-Protocol-(HTTP)-based, Representational-state-transfer (RESTful) application programming interface (API). Data objects exchanged with the server conform to the standard JavaScript Object Notation (JSON) format. Each object type in the encodeD system (or any like system built with SnoVault) can be accessed as an individual object or a collection (list) of objects. For example, to return a specific ENCODE experiment with the accession ENCSR107SLP, you can sent a “GET” request to the URL: https://www.encodeproject.org/experiments/ENCSR107SLP/. Accessions and unique names are aliased so that objects can be returned without referring to their object type in the URL. For example, https://www.encodeproject.org/ENCSR107SLP returns the same object. If the request is a browser, the JSON object returned is transformed into the webpage HTML by the ReactJS front end (Figure 1).

**Figure 1.**
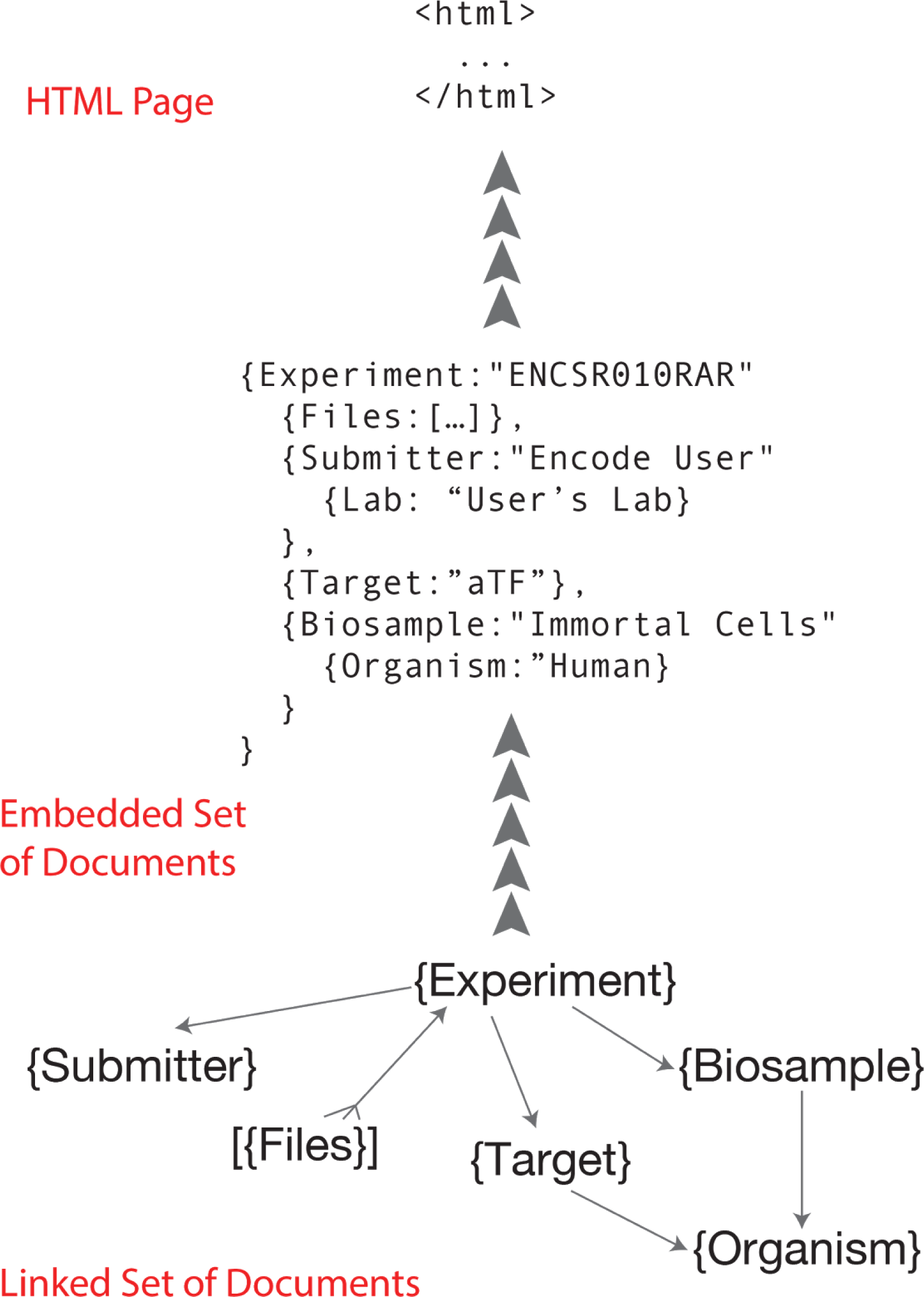
The normalized object graph (linked set of documents) stored in PostgreSQL is framed as an expanded JSON-LD document (embedded set of documents) before indexing into ElasticSearch and rendering in JavaScript.

Otherwise, Accept-headers (https://www.w3.org/Protocols/rfc2616/rfc2616-sec14.html) or the URL query parameter format=json can be used to request that the system return text JSON objects instead of HTML pages; but the URLs are precisely the same (Figures 2, 3). Because ElasticSearch indexes JSON objects, we can expose parts of the search API to allow command-line or programmatic access to our full facet search capability.

**FIgure 2.**
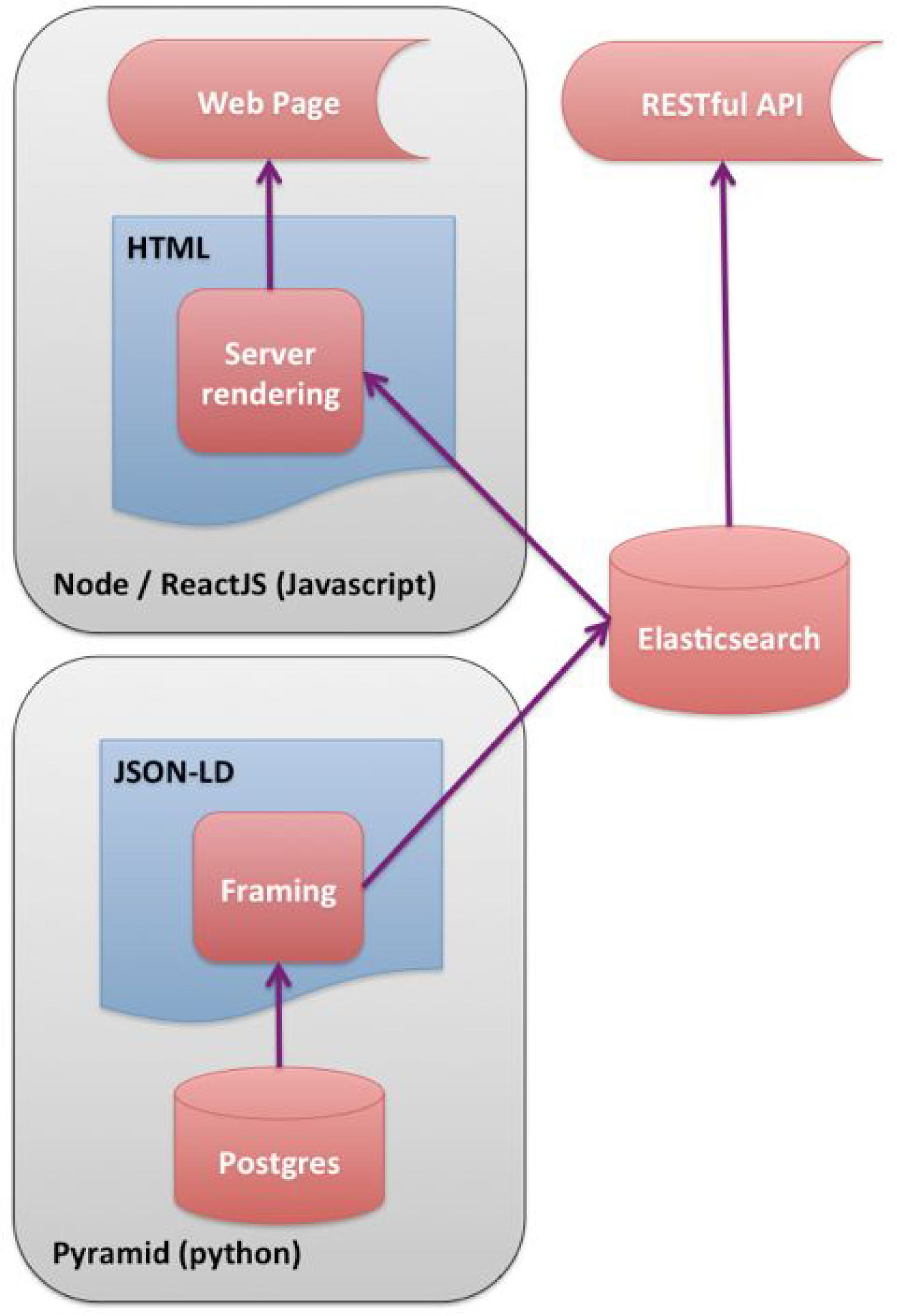
Schematic diagram of software stack showing different paths for page rendering. The most efficient rendering is with the HTML rendered on the server and the embedded documents indexed in ElasticSearch.

**Figure 3.**
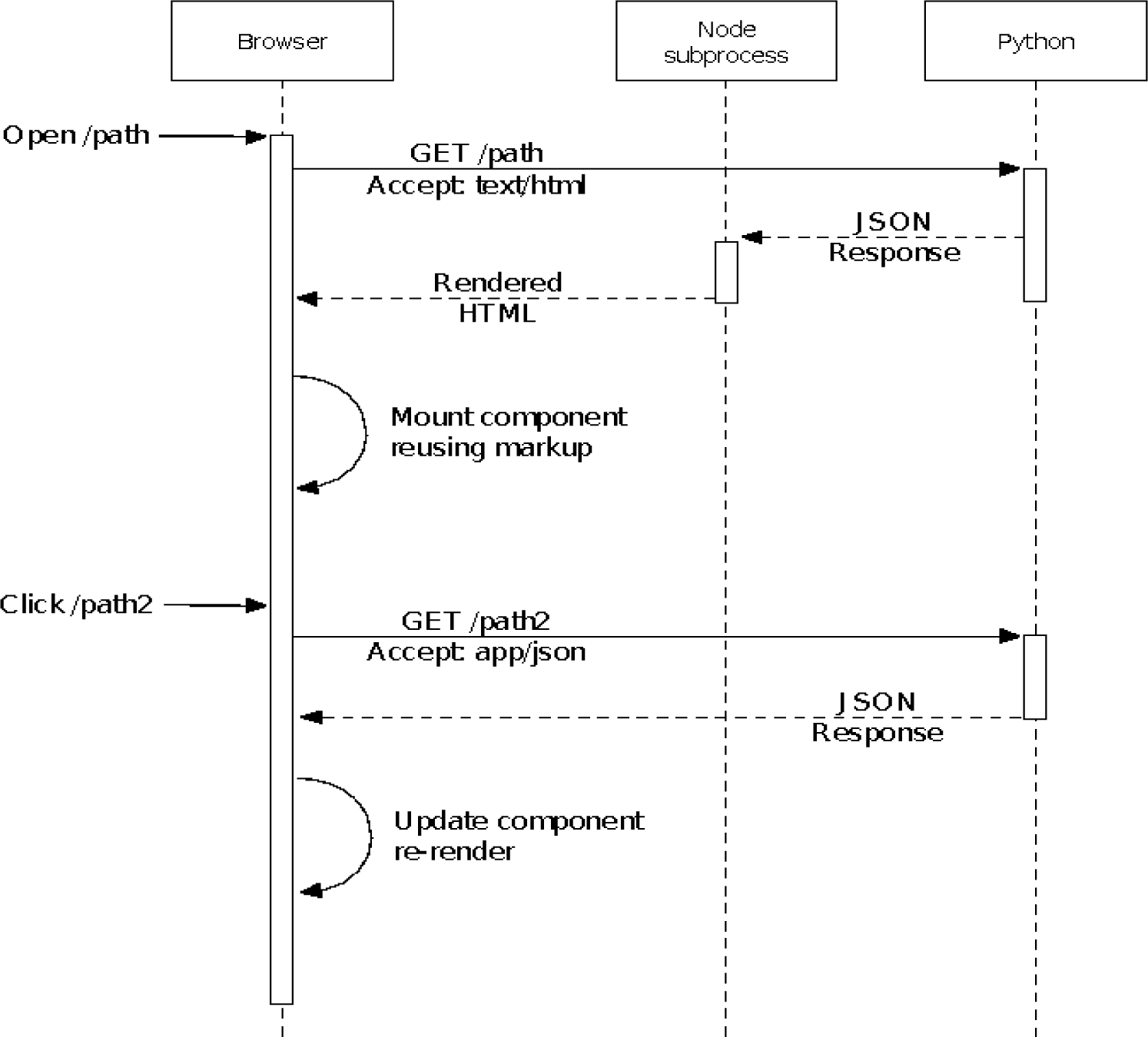
Initial page loads are rendered to HTML on the server for immediate display on the client. Once the JavaScript is fully loaded, subsequent page loads can be rendered on the client.

Members of the ENCODE consortium use either the REST API or HTML forms to submit data files and metadata to the DCC. New objects are created in the database by a POST request to the collection URL (for example, POSTing a valid biosample JSON object to the biosamples collection URL (https://www.encodeproject.org/biosamples/) creates a new biosample). Existing records are modified by a PATCH request to the object URL (for example, posting { “assay_term_name”: “RNA-seq”, “assay_term_id”: “OBI:0001271”} to https://www.encodeproject.org/ENCSR107SLP would change it from a ChIP-seq experiment to an RNA-seq experiment). Connection between experiments objects and biosample objects is maintained by the “linking” objects for Replicate and Library. A Replicate is assigned to one and only one Experiment, and references a specific Library created from (and linked to) the specific Biosample from which nucleic acid was extracted. Most submitters have several hundred experiments, and use simple python scripts to post their metadata and upload their datafiles to the DCC. For cases where a low throughput solution is expedient, a logged-in submitter can edit the properties of objects that belong to a lab they submit for, using a simple HTML form interface.

When new objects are added or edited, the objects are compared against the schema of the object to determine validity. A valid object (either POST, PUT, or PATCH) returns a “200” and an invalid one returns “422” with a brief description of the error. Our JSON Schema objects are, by design, quite permissive in the fields that they require. This approach was chosen to encourage ENCODE submitters to provide as much metadata as possible as early in the process, in effect “registering” experiments, biosamples, and antibodies before the complete metadata is marshalled and even before the experiments themselves have been completed. The earlier metadata is submitted to the DCC, the earlier we can catch irregularities, errors or data model confusions. We use PUT to create a wholly new object, POST to replace an existing object in its entirety (except for uuid), and PATCH to update only specific properties of an existing object.

One extension we have made to the JSON-SCHEMA representation of objects is to add what we call “calculated properties”. The JSON-SCHEMA definition for a particular object or document type is the minimum specification for properties that define the POSTing or creation of an instance of the object. For display and reporting purposes, the python code extends objects by properties that are calculated from one or more existing object properties or even linked properties. For example, this is how we show the files that belong to an experiment. In our system, files have a link to their “home” Dataset (of which Experiment is a sub-class). The “Files” of an Experiment are a property that appears in the return JSON objects but is not specified in the schema. Instead, they are calculated as a “reverse link” from the file objects themselves. Note that when calculated properties are indexed, updating a “child” object like File, it will result in the invalidation of the parent Experiment (or other home Dataset object) and trigger a re-indexing of the parent in ElasticSearch.

A similar feature we have added are something we call Audits. Audits were developed to act as cross-object type data integrity measures (in a way similar to triggers can be used by RDBS). Audits are small snippets of python code that are checked whenever an object added or updated (via POST, PUT, or PATCH). Because our system considers each object POSTed individually, we cannot use the JSON-SCHEMA representation to validate properties across linked data types. Generally speaking, a criteria of the object and its associated linked objects are compared to look for violations (e.g., human donor object should not have a “lifestage” property) or to track metadata inconsistencies (both replicates of an experiment should have the same biosample ontology term) or quality metric (bam files are typically expected to have have a minimum number of uniquely mapped reads). Audits are classified as ERROR (Red Flag), WARNING (Orange Flag), NON-COMPLAINT (Yellow Flag), or INTERNAL ACTION (Grey Ambulance) and are indexed with the page frame so that they can be displayed in the user interface (Figure 4). These audits help us retain a relational-like integrity across our sophisticated data graph, and give valuable feedback to submitter and wrangler alike. Finally, we use the audit system to report descrepancies between the ENCODE submission standards and quality control metrics and the true properties of the data and metadata. For example, if a particular experiment has low read mapped read depth (relative to standards set by the consortium for that particular class of experiment, it will be reported on the user interface and API as a “Warning” with a yellow flag.

**Figure 4.**
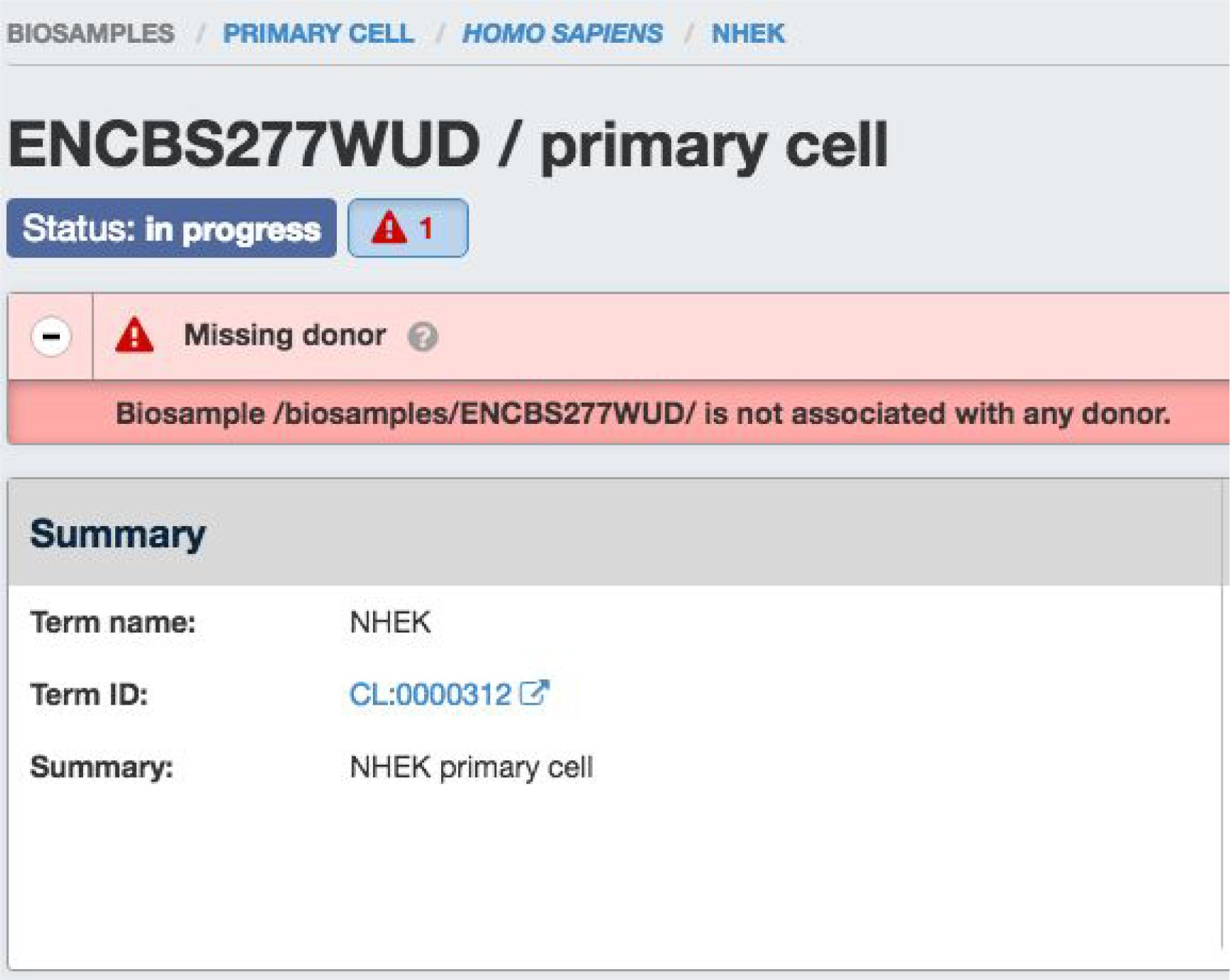
Example of the “missing donor” audit alerting submitters and DCC data wranglers that this epidermal keratinocyte biosample is missing the (human) donor object from whom it was derived.

One of the drawbacks of not using a fully relational database backend is that one cannot query with arbitrary joins across data types. For web applications, this is generally not a significant drawback, because each web page or API end point has to be specified in the source code anyway. Specifically, in our system we must either embed properties from linked object type B into object type A, or perform multiple queries (GET requests). There are some purposes for which this is inconvenient; for example creating database reports that span many objects. We outline here three different approaches that can be used if more arbitrary queries are needed. One approach is to simply use the python command line interpreter and the excellent requests module (http://docs.python-requests.org/en/master/) to download collections of objects in JSON and traverse them programmatically. This method is simple and straightforward but can be a little tedious if multiple queries across many different object types is wanted. A second approach is to query the postgreSQL database directly with psql. The objects are stored as JSON-type fields in postgreSQL, and starting with version 9.4 the individual properties of the JSON documents can be queried using the JSON operators implemented by postgreSQL. This method is not suitable if you are interested in calculated or embedded properties of the objects. Finally, we have added a utility to the SnoVault package that allows one to convert the JSON-LD object graph into Resource Descritive Format or RDF (https://www.w3.org/RDF/). A single API call to fetch all Objects (/search/?type=Item&limit=all&format=json) can be dumped to a file locally. Our script, jsonld_rdf.py uses rdflib (https://rdflib.readthedocs.org/en/stable/) converts objects to RDF in any standard format (xml, turtle, n3, trix and others). Once in RDF, it can be converted to a SPARQL (https://www.w3.org/TR/rdf-sparql-query/) store using a software package such as Virtuoso (http://semanticweb.org/wiki/Virtuoso.html). It could also be integrated with the EBI RDF Platform [32, 33]. In this way, the ENCODE metadata database (and any database created using SnoVault) can be made fully available to the semantic web.

## Front End

One of the advantages of using an object store where the documents are JSON objects, is that JSON (JavaScript Object Notation) is the native format used in JavaScript front end frameworks. No translations need to be made between the backend data structures and the front end. We have implemented the web front end to encodeD in ReactJS (http://facebook.github.io/react/)using JSX (http://jsx.github.io). ReactJS is a clean and efficient library for user interface programming in JavaScript, and JSX allows us to use XML-like stanzas in place of an HTML or XML templating system. We further optimized the performance of ReactJS by compiling (from JavaScript to XHTML) the pages on the server using a node (https://nodeis.org) process (Figures 2 and 3). Most object types have useful landing pages in HTML that provide a human-friendly version of the raw JSON data, while collections of objects are accessed via the faceted search interface (https://www.encodeproject.org/search/ and Figure 6A). We configure the facets for each object type in the JSON schema objects. These are then translated into ElasticSearch aggregations providing rapid filtering and winnowing through data. We have recently implemented an Experiment matrix view (https://www.encodeproiect.org/matrix/?tvpe=Experiment and Figure 5) and a spreadsheet report view ( https://www.encodeproiect.org/report/?tvpe=Experiment Figure 6) within the search, expanding the modes in which our users can process our data. Individual experiments or sets of experiments with browser-viewable files can be directly viewed at the UCSC genome browser[34]via a track hu b[32]created on the fly (Visualize Button), and we provide a metadata driven method for downloading all data files associated with a specific search using the “Download” button. From searching, one can drill down to a specific object page to get detailed information on all the metadata collected for an object.

**Figure 5.**
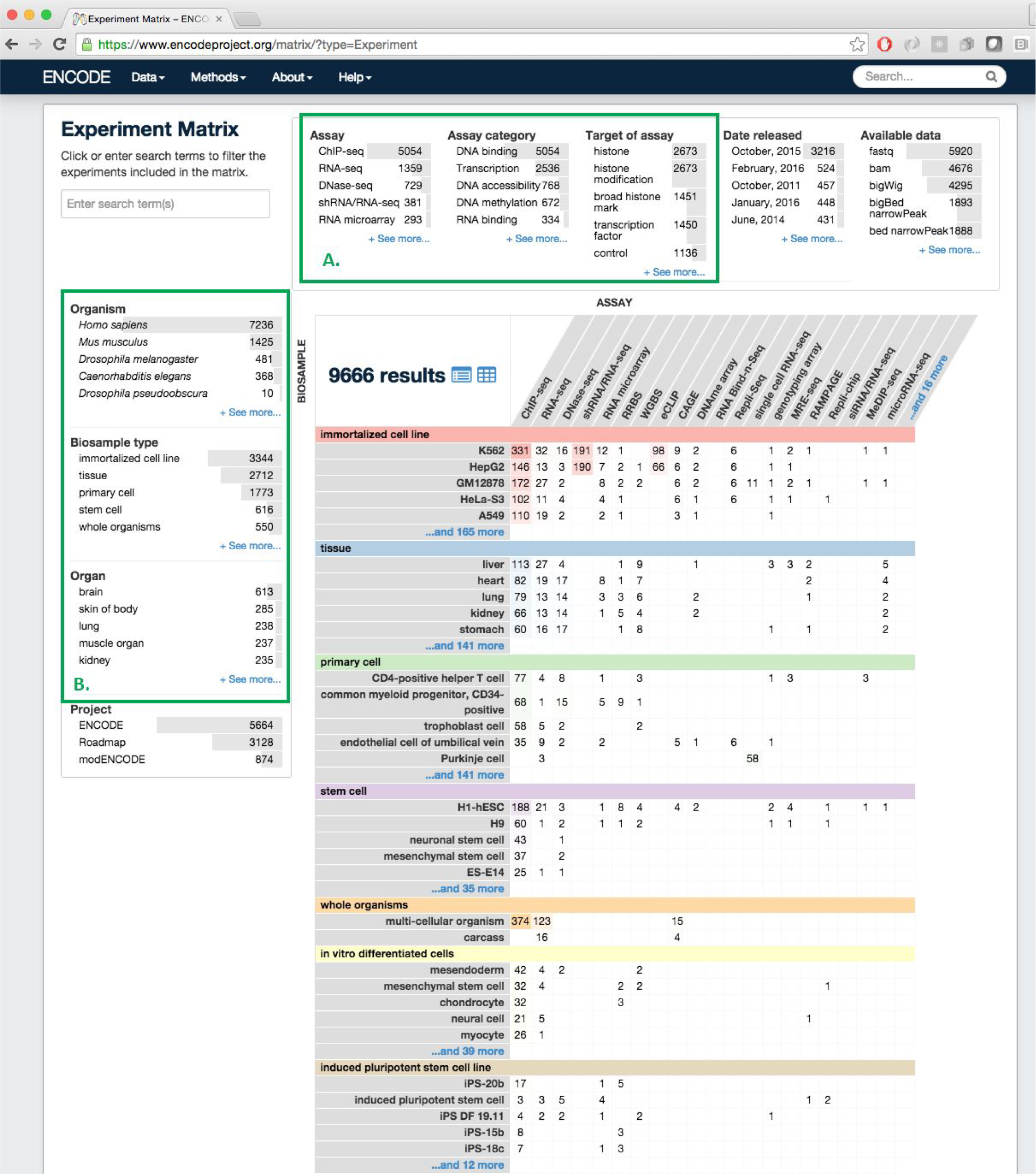
ENCODE Portal Data Matrix which shows all experiments released at the ENCODE Portal, including the Roadmap for Epigenomic Mapping Consortium (REMC). Experiments are organized by their biosample (tissue, cell or cell line) on the Y axis and by Assay type on the X-axis **(A)** Facets select specific properties, such as target (histone, transcription factor) or experimental type (ChIP-seq, RNA-seq, etc.) **(B)**Facets apply specifically to the biosample, including organism (human, mouse, fly, worm), type (tissue, immortalized cell line, stem cell, etc.) or organ system (as inferred from ontological relations).

**Figure 6.**
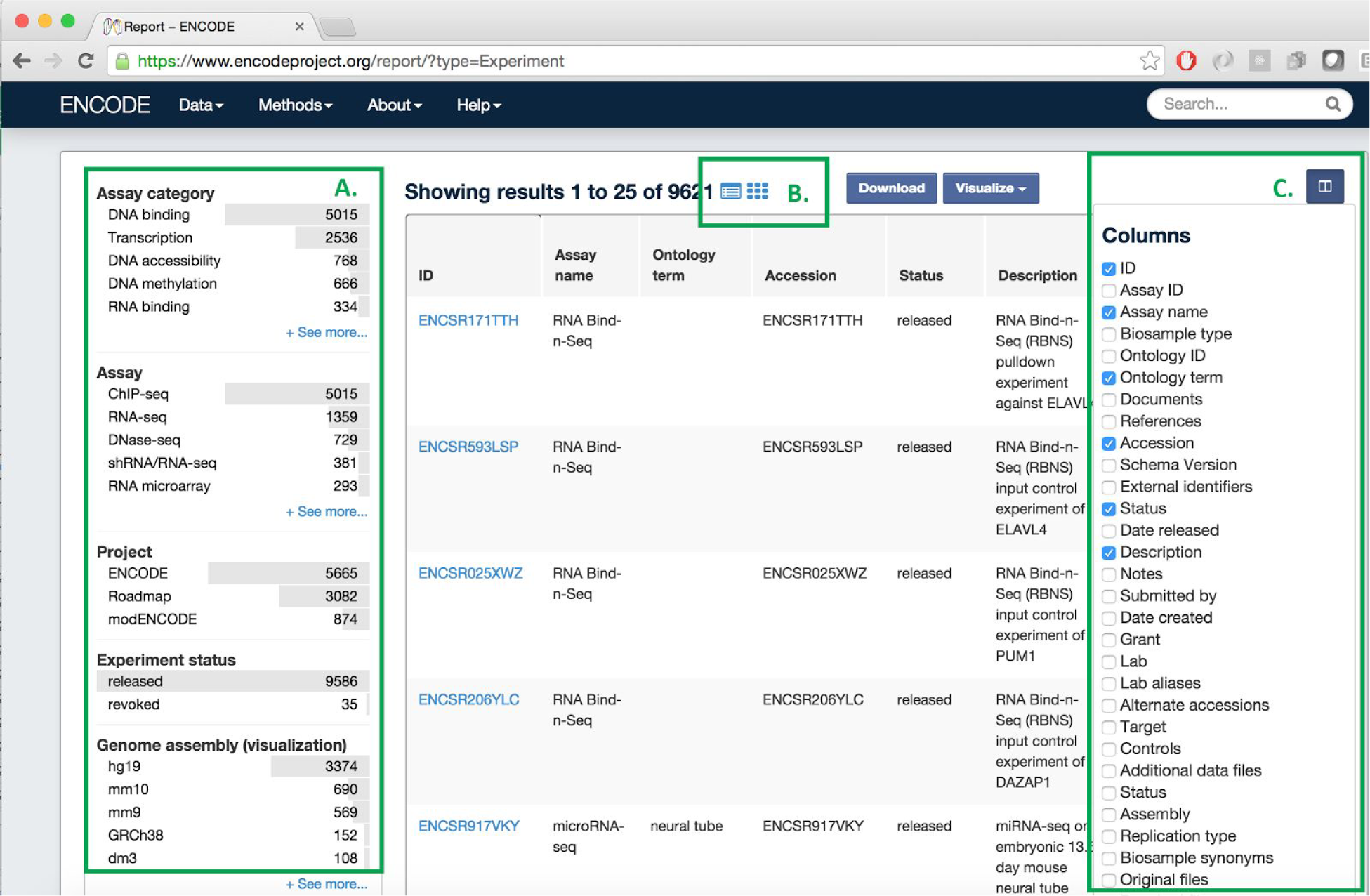
ENCODE Portal Report page, where metadata can be downloaded in spreadsheet format (csv). Report views exist for all collection searches, for example, Experiment, Biosample or Antibody. **A**. Standard ENCODE Assay facets to filter the rows of interest (in this case Experiments). **B**. Toggle between the report, matrix, and standard search output. **C**. Select the columns (individual properties) that will appear in the columns of your spreadsheet.

## Using Github and the Cloud for Rapid Development and Deployment

To enable rapid software development and robust deployment, the ENCODE DCC has implemented cloud dev-ops (development operations) code directly into the encodeD system. Using the cloud effectively is a combination of a few small python and well-defined coding practices. Our coding practices start with git, and github. Git is a open-source distributed version control software that is the state of the art in current software engineering. We manage bug requests and feature requests (”Issues”) in tracking software, where each issue is assigned a number and a developer. The developer creates a git branch specific to this issue and works on his or her local version of the software (including a small text database or “fixture”). The branch code is pushed daily to github so that the team is always kept up to date. Github is a web-based Git repository hosting service, which is free to open-source projects. We have written hooks to a web-based continuous integration service called travis-ci (http://travis-ci.com), such that when any branch is pushed to github, our testing suite is initiated on a virtual machine hosted by travis-ci. This service is also free for open-source projects. Every 2-3 weeks, completed features are collected into a “release candidate”, merged together, put through a manual Quality Assurance (QA) process, and released to production. Release candidates or even specific feature branches that require full production databases or deeper QA, can be installed on specific demonstration machines via a script that creates and “names” a virtual machine in the Amazon Cloud (AWS).

We use postgreSQL Write-Ahead Logging

(WAL;http://www.postgresql.org/docs/9.4/static/wal-intro.html) and the sotware package WAL-E (https://github.com/wal-e/wal-e) to maintain a consistent database across our production, test, and various “feature-demo” instances created by engineers and data wranglers for testing and QA. A simple flag to the deploy script configures each instance to be a production candidate, test (global testing) or demo (local, in-house testing machine), and they immediately create a local copy of the postgreSQL database via the WAL logs. The difference between a production instance and any other instance, is that the production instance acts as the postgreSQL “master” and is the only machine authorized to *write* to the WAL logs or to upload files to the production Amazon S3 bucket.

## Using encodeD or SnoVault In Your Project

### Open Source Repository

All of the source code created by the ENCODE DCC is available from GitHub: http://github.com/ENCODE-DCC/. SnoVault is a completely generic hybrid object database with ElasticSearch interface. It is completely data agnostic and could be used for any web database purpose. Our system used for the ENCODE DCC metadata database is at http://github.com/ENCODE-DCC/encoded. which could be adapted to other projects that specific store genomic data and metadata

### PyPi Package

We have extracted the main Pyramid/JSON-SCHEMA/JSON-LD/ElasticSearch framework to a separate repository, and created a python package called SnoVault (http://github.com/ENCODE-DCC/snovault). SnoVault is available from PyPi or other python distribution systems. The main object-relational backend can be used for any project regardless of content. Creating application specific code starts with modeling your data in JSON using JSON-SCHEMA and JSON-LD. A small amount of custom code is required to get the bare bones application running for item and collection pages, and from there frontend components using ReactJS or any other system can be used to give your application a unique user experience. We have used the encodeD system to create a curation application and portal for the ClinGen project (http://clinicalgenome.org: http://github.com/ClinGen/clincoded), and it is currently being evaluated by other genomiomics Data Coordination Centers at Washington Universty (Saint Louis) and Harvard Medical School.

## Summary

When the ENCODE DCC project moved to Stanford University in 2012, we had to develop a software system that could handle all existing data and metadata from ENCODE, incorporate distinct and additional metadata from modENCODE and REMC, as well as all future ENCODE metadata. In many cases, the experimental assays, reagents, and protocols were not yet fully defined when we began our work on encodeD, while experimental data was being continuously created from existing ENCODE production labs. In short we had to develop a system flexible enough to handle nearly arbitrary experimental definitions within the field of genomics and epigenomics, yet still maintain strong data integrity and control the input specification to preserve univocity in our data and metadata descriptions. Our schema changes with nearly every software release, and the software is released in place every 3-4 weeks, with almost zero disruption to submitting or viewing users.

## Acknowledgements

We wish to thank all participants in the ENCODE Consortium for our collaborations that have enhanced the metadata definitions, as well as ENCODE Portal users for their useful feedback.

